# Functional Recovery from Human Induced Pluripotent Stem Cell-Derived Dopamine Neuron Grafts is Dependent on Neurite Outgrowth

**DOI:** 10.1101/2022.04.19.488213

**Authors:** Rachel Hills, Jim A. Mossman, Andres M. Bratt-Leal, Ha Tran, Roy M. Williams, David G. Stouffer, Irina V. Sokolova, Pietro P. Sanna, Jeanne F. Loring, Mariah J. Lelos

## Abstract

Transplantation of human pluripotent stem cell-derived dopaminergic (DA) neurons is an emerging therapeutic strategy for Parkinson’s disease (PD). In this study, we demonstrate, for the first time, that that functional recovery after DA neuron transplant is critically dependent on graft-host integration, and not dependent on graft size or DA neuron content. Specifically, in anticipation of an autologous DA neuron transplant strategy, we studied two human induced pluripotent stem cell lines derived from people with idiopathic PD. We confirmed the cells’ ability to differentiate into mature DA neurons in vitro by assessing electrophysiology and depolarization-induced neurotransmitter release. To evaluate efficacy, we transplanted DA neuron precursors into a hemiparkinsonian rat model. We differentiated the cell lines for either 18 (d18) or 25 days (d25) to investigate the effect of cell maturity on efficacy. We found comparable graft survival in d18 and d25 groups from both cell lines, but behavioral analysis revealed that only d18 preparations, from both cell lines, resulted in recovery of motor impairments. Immunohistochemical analysis did not reveal any DA neuron-associated markers that correlated with efficacy. However, we found striking and consistent differences in graft neurite outgrowth between the two culture timepoints. The functional grafts from d18 cells had more outgrowth than non-functional d25 grafts. A time course of gene expression during differentiation of the cell lines revealed differences in genes associated with DA neuron development and neurite plasticity. These results are the first to demonstrate that graft-host integration, and not DA neuron content, is key to graft-induced functional recovery. The gene expression profiling may offer insight into the optimal developmental stage for graft efficacy.

## Introduction

Therapeutic interventions that alleviate motor symptoms in Parkinson’s disease (PD) typically target replacement of lost dopamine (DA) transmission within the basal ganglia in order to overcome the impact of nigrostriatal degeneration (Henchcliffe and Parmar, 2018). Through ectopic transplantation, cell replacement therapies seek to restore lost connections of DA neurons to medium spiny neurons in the striatum. Clinical trials using fetal DA cells as a reparative strategy for PD have demonstrated long-term recovery of function in some patients (Barker *et al*., 2013) and graft survival for over 20 years (Li *et al*., 2016).

Human pluripotent stem cells (hPSCs: human embryonic stem cells or human induced pluripotent stem cells) can be differentiated into ventral midbrain DA (vmDA) cells and transplanted, and these cell therapy products have been widely reported to survive, integrate, release DA and alleviate functional impairments in rodent models of PD (Kriks *et al*., 2011; Kirkeby *et al*., 2012; Grealish *et al*., 2015; Lelos *et al*., 2016). Cell therapies for PD are moving into an exciting era, with several first-in-human clinical trials using hPSC derived neural precursors recently begun, or due to commence (Barker *et al*., 2017; Kim, Koo and Studer, 2020; Schweitzer *et al*., 2020; Takahashi, 2020).

Most current strategies for cell replacement have relied on a single hPSC line to generate DA neuron precursors for transplantation. Since these allogenic approaches require immunosuppression and its associated risks, we are investigating the feasibility of autologous DA neuron replacement. In this study we used induced pluripotent stem cells (iPSCs) from two individuals with idiopathic PD. The primary aims were to assess the reproducibility of the generation of transplant-ready DA neuron precursors, and to identify the point during *in vitro* differentiation for optimal integration and efficacy of the iPSC-DA grafts. We assessed DA precursors from each cell line at two different stages of differentiation by transplantation to a hemiparkinsonian rat model. We chose 18 days (d18) and 25 days (d25) of differentiation *in vitro*, and to better understand the composition of the cultures, we phenotyped the two stages with transcriptome profiling. Our data show that regardless of cell line or differentiation stage, all transplanted preparations survived and matured into grafts rich in vmDA neurons. However, we found that there was a striking difference in efficacy between d18 and d25-differentiated cells for derivatives of both iPSC lines. The earlier stage precursors consistently alleviated motor impairments in the rat model as measured six months after transplantation, while the more mature d25 cells consistently failed to reverse motor symptoms in the model. Immunohistochemical analysis of the iPSC-DA grafts suggested that classic measures of graft volume or DA neuron content did not distinguish efficacious from non-efficacious transplants. Instead, significant differences in the extent of neurite outgrowth from the grafts was the factor that correlated with the ability to elicit behavioral recovery. Transcriptome analysis for cell culture phenotype revealed differences between efficacious and non-efficacious DA precursors in expression of genes related to the stage of neuronal development. The results of this study show that differentiation is reproducible in two different patient-derived iPSC lines, and that the developmental stage of the cultured cells determines their ability to reverse motor deficits in a rat model of PD, which is dependent on their ability to extend neurites into the host brain.

## Results

Two iPSC lines (410, 411) were generated from fibroblasts harvested from two people with idiopathic PD (**Figure 1**). These two iPSC lines were differentiated *in vitro* to DA neuron precursors for 18 or 25 days to investigate the optimal stage for transplantation. To confirm that the precursors were capable of differentiation into mature DA neurons, both cell lines were differentiated for a further 12 weeks *in vitro* to mature DA neurons. The precursors harvested after 18 days (410-d18; 411-d18) or 25 days (410-d25; 411-d25) of differentiation were used for transplantation into a rodent model of PD. At the same stages, replicate cultures were analyzed by mRNA sequencing to examine their phenotypic characteristics.

**Figure 1.**
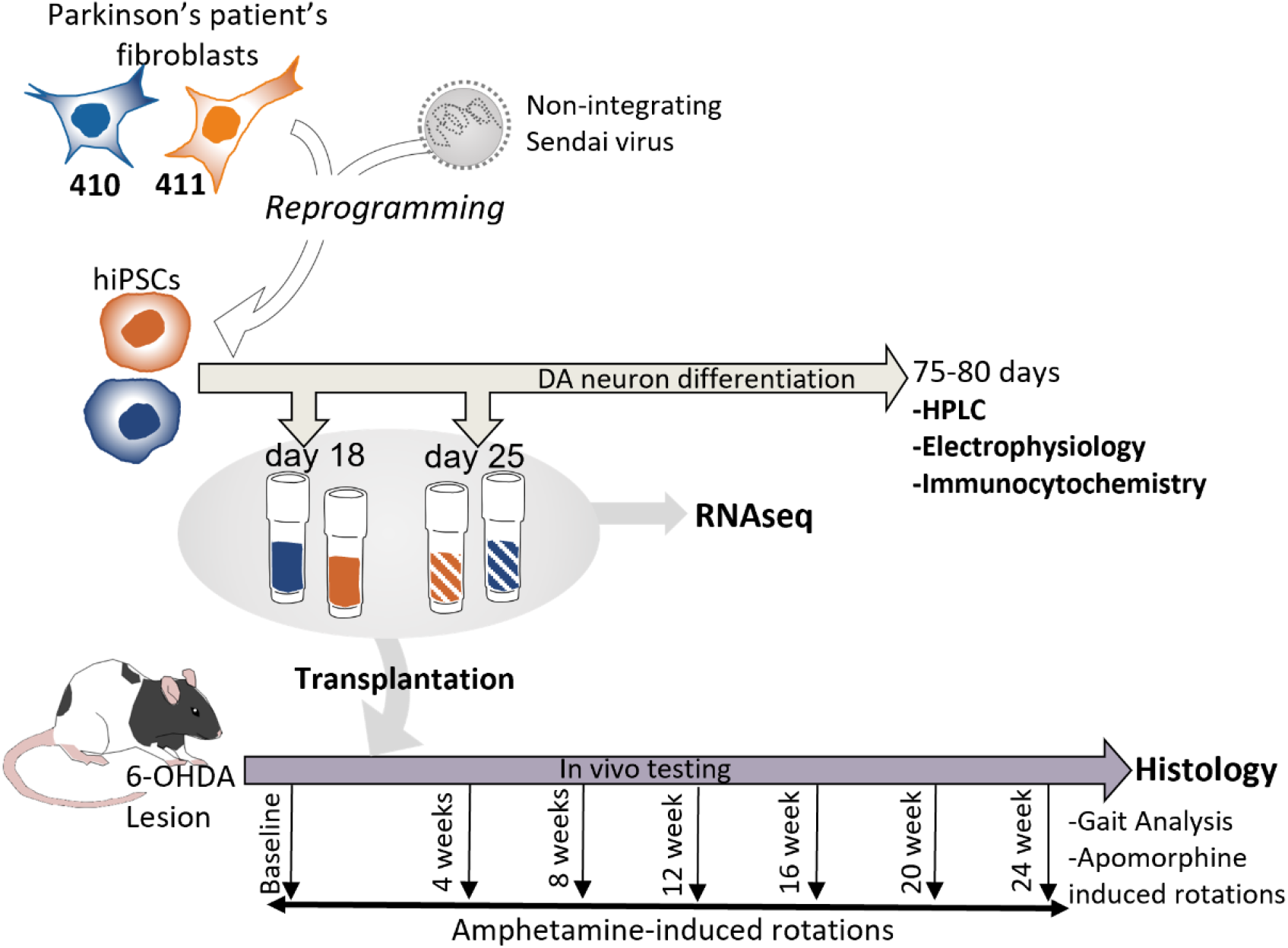
Study design showing experimental progression from PD donors’ fibroblasts to CTPs, and the main outcome measures from both *in vitro* and *in vivo* investigation of their suitability as a dopamine cell replacement source.

Both cell lines were transplanted into the 6-OHDA lesioned hemiparkinsonian model of PD, at both stages of the differentiation protocol. Other groups of rats remained as naïve controls or lesioned controls. Rats underwent behavioral testing pre- and post-transplantation and brains were taken for histological analysis at 24 weeks post-graft. Histological analysis of the grafts included markers of human cells (HuNu) and mature DA neurons (TH, GIRK2, AADC), as well as two methods of analyzing the extent of neurite outgrowth from the TH+ neurons.

### Both iPSC lines differentiated into mature dopamine neurons in vitro

To validate their ability to differentiate into dopamine neurons, iPSCs were cultured for approximately 12 weeks in maturation conditions. Both 410 and 411 cell lines differentiated into mature DA neurons, as evidenced by their neuronal morphology and expression of tyrosine hydroxylase (TH) (**Figure 2A**). Furthermore, electrophysiological analysis showed that they had spontaneous spiking activity typical of A9 substantia nigra neurons (**Figure 2B-D**). When depolarized, the neurons released dopamine (**Figure 2E**).

**Figure 2.**
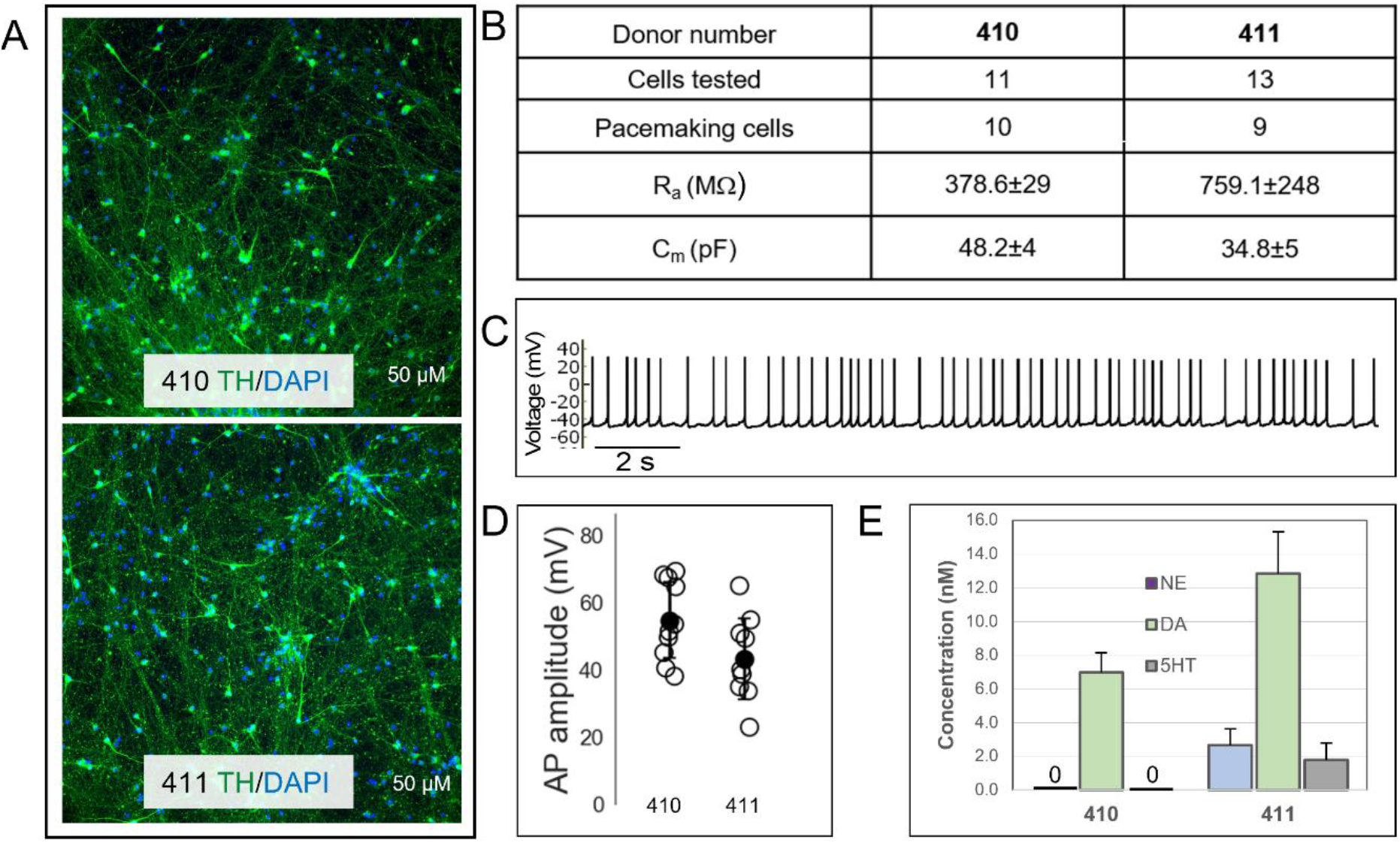
Characterization of dopaminergic neuron differentiation and maturation from two PD donor iPSC lines. (A) iPSCs from 410 and 411 cell lines that were differentiated to day 80 *in vitro* express dopamine neuron marker tyrosine hydroxylase (TH; green), (B) Single-cell patch clamp analysis of day 75 cells showed that most of the neurons had pace making activity and mature membrane resistance (Ra) and capacitance (Cm). (C & D) Action potential amplitudes were maintained close to 60mV. (E) HPLC analysis of day 88-89 cultures stimulated with KCl shows release of dopamine (DA) and small amounts of norepinephrine (NE) and serotonin (5-HT).

### Behavioral analysis revealed motor recovery after transplantation of earlier (d18) but not later (d25) DA neuron precursors

The amphetamine-induced rotation test detects DA imbalance within the basal ganglia in lesioned rats; systemic injection of amphetamine causes DA release from the unlesioned side rats and evokes ipsilateral rotations. As expected, control rats did not rotate at any timepoint in response to amphetamine, while lesioned rats consistently rotated in response to amphetamine. The grafts of 410-d18 and 411-d18 DA precursors restored the rotational bias over 24 weeks. In contrast, the 410-d25 and 411-d25 graft groups did not recover from the imbalance caused by the lesion (**Figure 3A-B**; Time*Group: F_30,270_= 5.95, p≤ 0.001, see **Supplemental Figure 1** for statistical data table).

**Figure 3.**
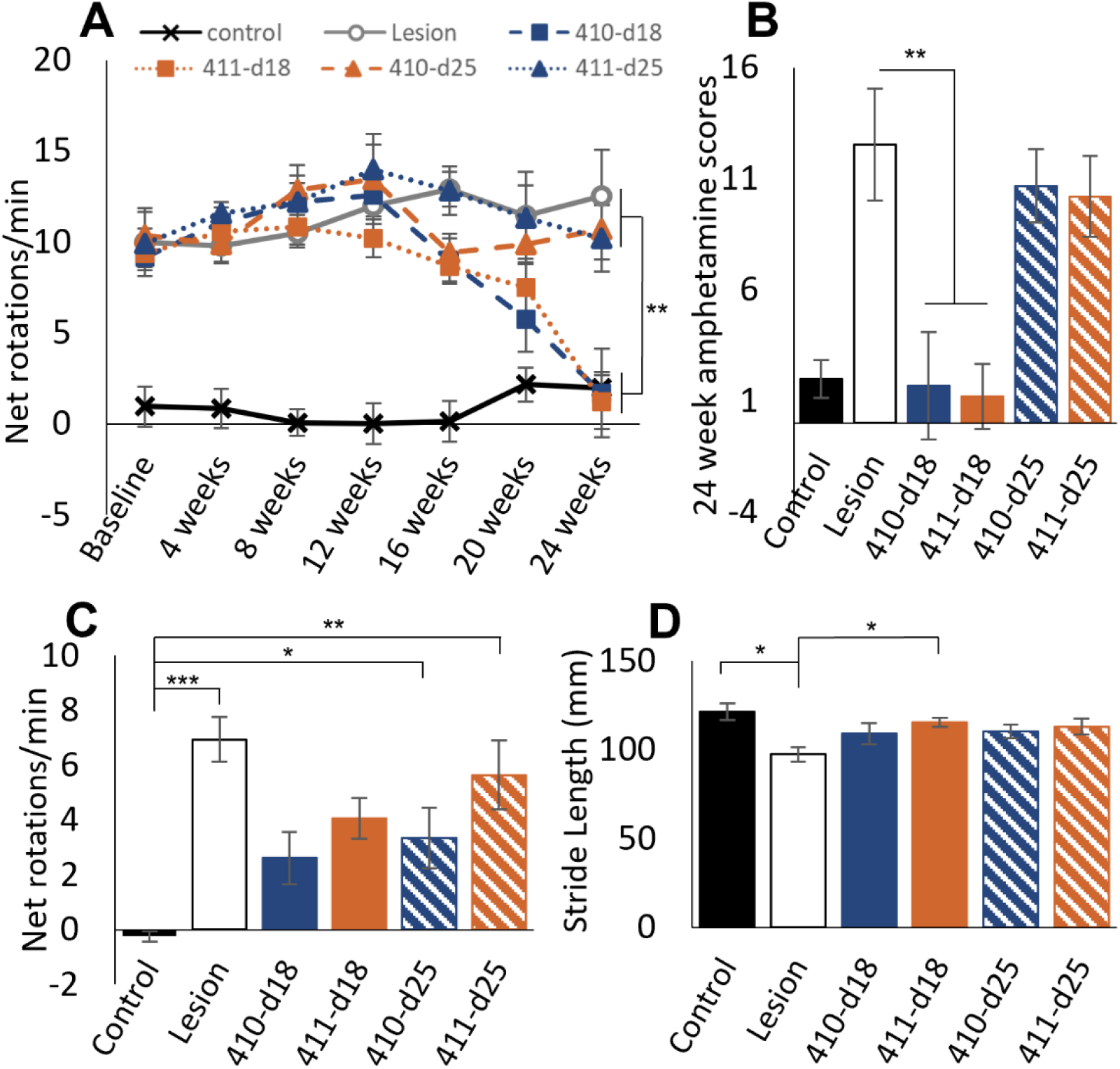
Effect of iPSC-derived cells on motor function. Raw data showing performance of Control, Lesion, 410-d18, 411-d18, 410-d25 and 411-25 grafted rats on amphetamine-induced rotation test across the full test period (A), and net rotation score at 24 weeks post-graft only (B). Data are presented for the apomorphine-induced rotation test as net rotation score at 24 weeks post-graft (C), and stride length on gait analysis at 24 weeks post-graft (D). *p≤ 0.05, **p≤0.001. Error bars = SEM.

For the apomorphine-induced rotation test, systemic injection of the dopamine receptor agonist, apomorphine, induces contralateral rotation in 6-OHDA-lesioned rats, caused by super-sensitivity of the receptors in the lesioned hemisphere. At 24 weeks post-graft, rats in the lesion-only group (p<0.001) and rats with 410-d25 and 411-d25 grafts (p<0.05) rotated significantly more than the control group in response to apomorphine. However, rats grafted with cells from the earlier stage of differentiation (410-d18 and 411-d18) did not differ from unlesioned control rats, indicating that improvement in the rotational bias was conferred by both cell lines differentiated to d18 (**Figure 3C**; Group: F_5,45_= 6.34, p<0.001, see **Supplemental Figure 1** for statistical data table).

Gait analysis, which is sensitive to changes in stride length and posture caused by DA loss in DA lesioned rats, was used as a non-drug-induced test of motor function. Lesioned ungrafted rats had a decreased stride length compared to naïve control group animals (**Figure 3D**; F_5,45_= 3.27, p=0.013; lesion vs control p=0.007). In contrast, rats grafted with cells from the earlier differentiation stage showed improvement in this motor feature, with increased stride length relative to lesioned controls (p=0.047; see **Supplemental Figure 1** for statistical data table).

In summary, by multiple behavioral analyses, the hemiparkinsonian rats transplanted with the earlier (d18) stage precursors showed reversal of deficits, while those transplanted with the later (d25) stage precursors did not recover.

### Differences in graft size or DA neuron content do not explain the difference in efficacy between d18 and d25 precursor grafts

The simplest explanation for the failure of the later stage (d25) precursors would be that the cells didn’t engraft or formed very small grafts with few DA neurons. This was not the case. Immunohistological analysis was used to quantify the number of engrafted human cells (HuNu+), and cells positive for mature DA neurons (TH+, AADC+, GIRK2+) in grafts. The analysis revealed variations dependent on cell line (“Cell line”) and/or days *in vitro* (“DIV”); however, these characteristics could not explain the difference in functional efficacy between the d18 and d25 grafts. **Figure 4** shows TH immunolabelling of representative examples of d18 and d25 grafts at 24 weeks post-transplant. In general, d18 grafts were larger (**Figure 5A,B**) and had more TH+ (**Figure 5D**), GIRK2+ (**Figure 5G,E**) and AADC+ cells (**Figure 5J**) [DIV, minimum F_1,11_= 6.02, p<0.02]. The density of TH+ neurons (**Figure 5C**) and the percentage of GIRK2+ neurons (**Figure 5H**) was also greater in d18 grafts [DIV, minimum F_1,11_= 6.32, p<0.05]. But comparison of similar sized grafts makes clear that none of those factors was critical to the behavioral outcome. This is illustrated in **Figure 5** by comparing a group of grafts of similar size but with different behavioral outcomes. Day 25 (410-d25) and day 18 (411-d18) grafts had different behavioral outcomes, but the non-efficacious day 25 grafts were slightly larger (**Figure 5A,B**), had comparable TH+ density (**Figure 5C**), and contained more TH+ (**Figure 5D**), GIRK2+ (**Figure 5G**) and AADC+ neurons (**Figure 5J**) than the day 18 grafts [Cell Line, minimum F_1,11_= 6.97, p<0.02]. In summary, behavioral recovery did not correlate with the number of HuNu+ cells, graft volume, total number of TH+ cells, AADC+ cells, GIRK2+ cells, nor density of TH+ cells (**Figure 6F**).

**Figure 4.**
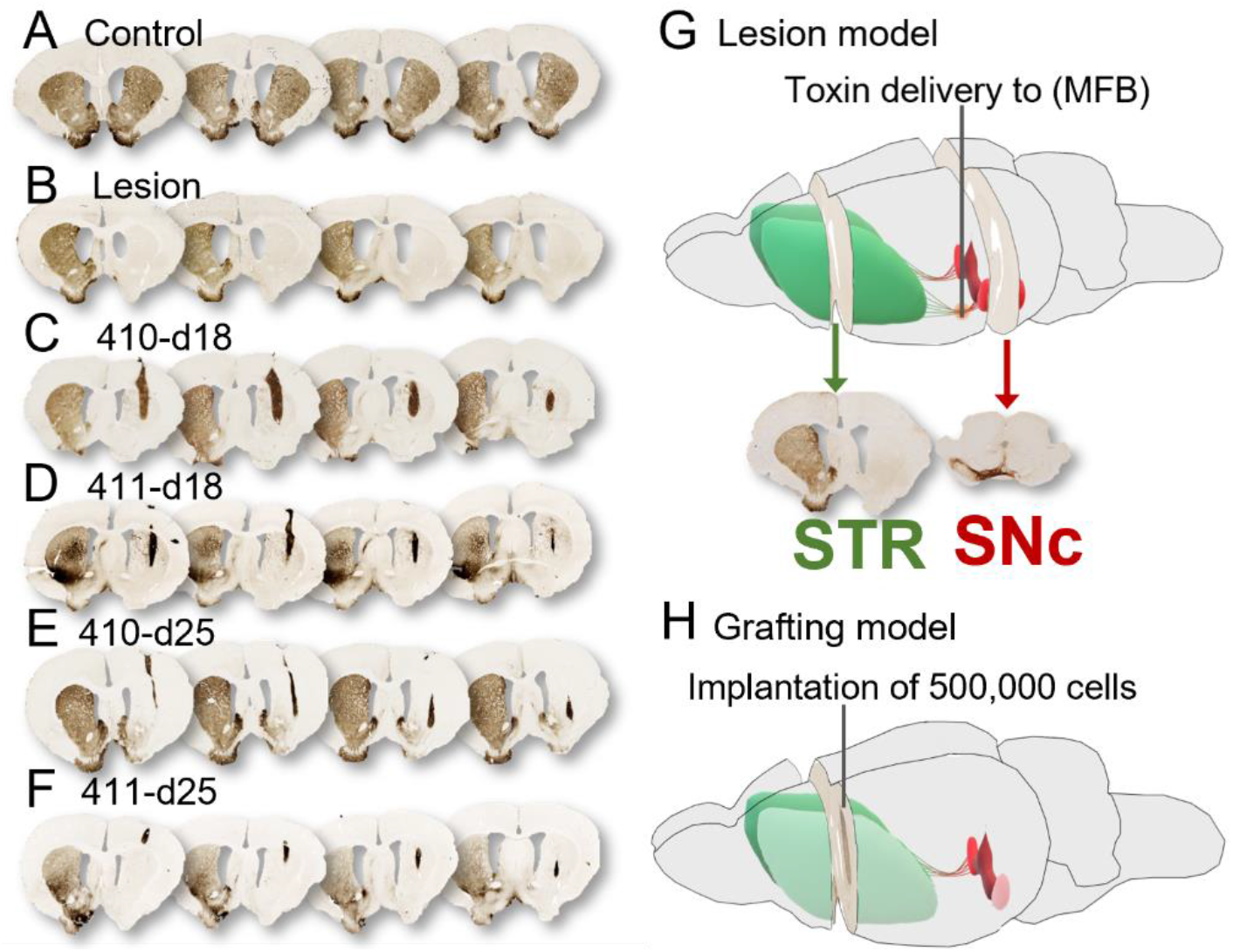
Representative examples of immunolabelling of tyrosine hydroxylase-positive (TH+) neurons in iPSC-DA CTPs in the neostriatum of 6-OHDA lesioned rats. (A) naïve controls, (B) 6-OHDA lesion controls, (C) 410-d18 grafts, (D) 411-d18 grafts, (E) 410-d25 grafts and (F) 411-d25 grafts. Schematics showing unilateral 6-OHDA medial forebrain bundle (MFB) lesion model with accompanying unilateral TH depletion in immunolabelled brain sections in the striatum (STR) and substantia nigra pars compacta (SNc) and cell replacement strategy of grafting surgery (H).

**Figure 5.**
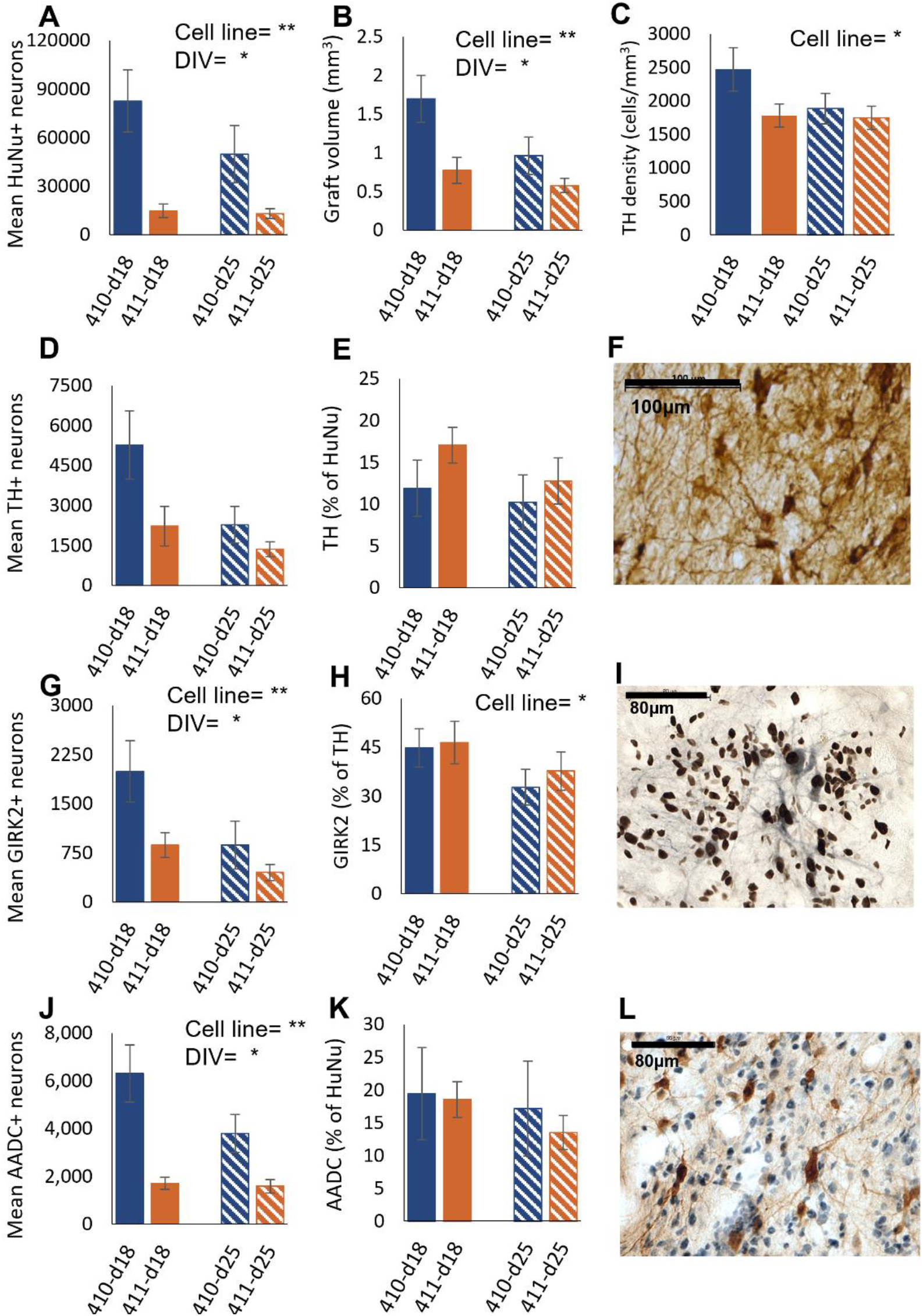
Immunohistochemical analysis of d18 and d25 iPSC-derived grafts. Based on histological analysis of the grafts from d18 and d25 preparations, we present the total mean HuNu+cells (A), graft volume (B), the density of TH+ cells per mm^3^ (C), total TH+ neurons (D), percentage of TH+ cells out of total HuNu+ cells (E), the total GIRK2+ cells (G) and the percentage of GIRK2+ cells out of TH+ cells (H). The total AADC+ cell data are depicted in (J) and the percentage of AADC+ cells relative to HuNu+ cells is in (K). Representative immunohistochemistry is presented for TH (F), GIRK2 (blue) and HuNu (brown) in (I) and AADC (brown) and HuNu (blue) in (L). Main effects of cell line or days in vitro (DIV) are stated, with p*≤0.05, p**≤0.001, error bars=±SEM.

**Figure 6.**
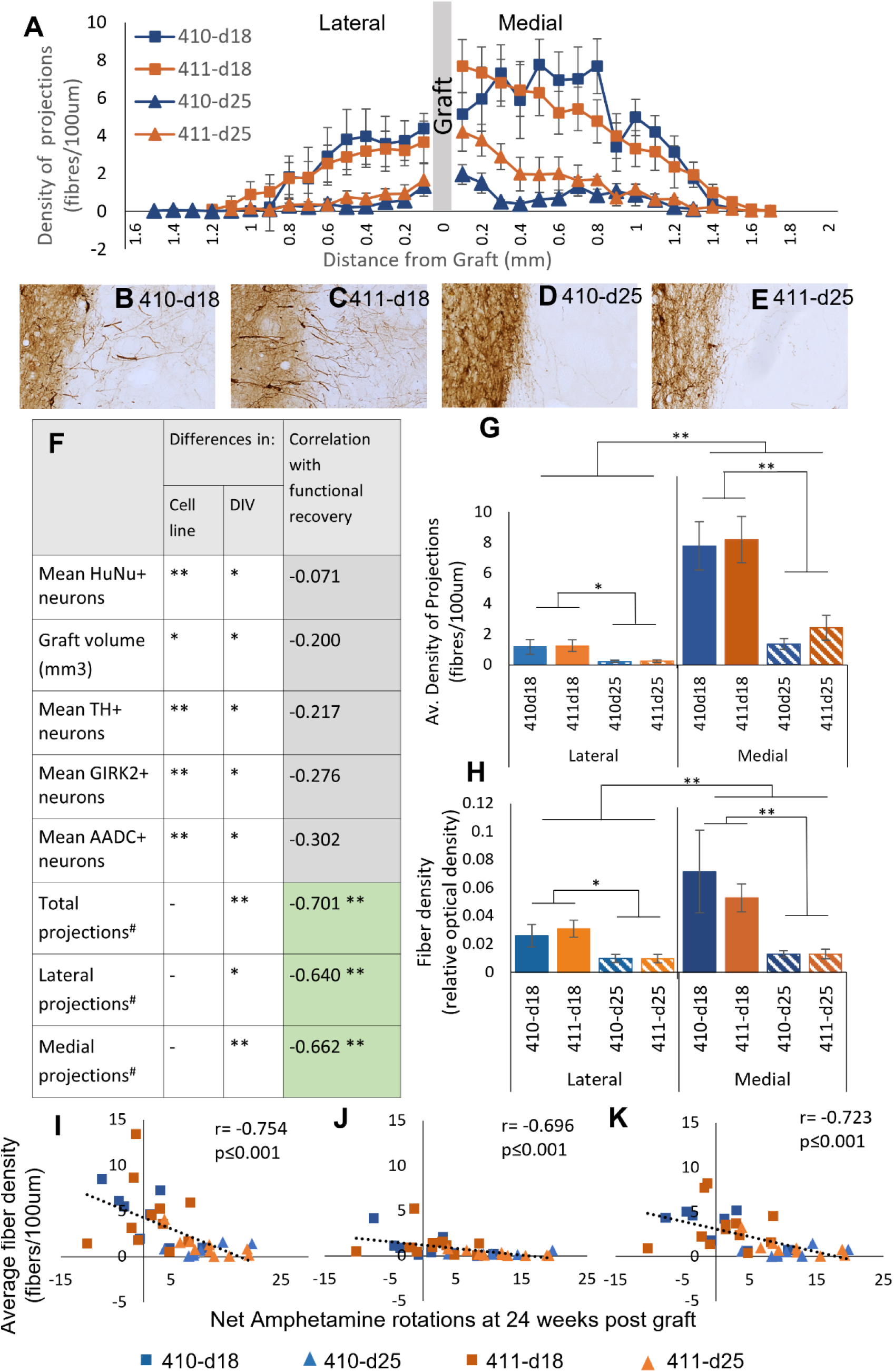
Stereological analysis of TH+ neurite outgrowth. The mean density of neurite projections (per 100μm) is depicted as distance from the graft in both a lateral and medial direction for each graft group (A). (B-E) Representative images of the graft/host border demonstrating increased neurite outgrowth from the graft core in both day 18 groups (B and C) compared to relatively little neurite outgrowth from day 25 grafts (D and E). Manual fiber counting demonstrated that the average density of projections was higher in both 410-d18 and 411-d18 grafts (G) and this was further supported by an objective measure of fiber density (optical density) in the striatum adjacent to the graft (H). The projection data are shown split into projections into the lateral or medial striatum, with more medial innervation evident overall (G and H). Net amphetamine-induced rotation score at 24 weeks post-graft correlated significantly with overall density of graft projections (I), as well as both lateral (J) and medial (K) projection densities independently (J). Neurite outgrowth was the histological variable that correlated with behavioral improvements (F).

Representative histology for TH+, GIRK2+, AADC+ and HuNu+ neurons is depicted in **Figure 5F,I,L**.

### Functional improvements correlate with significantly more extensive neurite outgrowth from grafts of earlier (d18) DA precursors

Manual counting of fibers revealed 4-fold higher average density of projections from d18 grafts compared to d25 grafts (**Figure 6G**; DIV. F_1,11_= 45.73, p<0.001). Significantly more neurite outgrowth was also evident in d18 grafts when the tissue was analyzed using unbiased optical density estimates via computerized image analysis (Image-J) (**Figure 6H**; DIV. F_1,11_=14.31, p=0.003). As the distribution of projections was not homogenous across the striatum, the y-intercept of projection trend lines was also analyzed, and again confirmed significantly more neurite outgrowth from d18 grafts (**Supplemental Figure 2A**; DIV: F_1,11_= 44.16, p<0.001). Finally, to eliminate the influence of any differences in graft size, projection data were normalized to the number of TH+ cells within the graft; there were still significantly more projections emanating from d18 grafts (**Supplemental Figure 2B**; DIV: F_1,11_= 18.07, p≤0.001). Together, these analyses suggest that differences in graft integration are due to underlying features of the transplanted cells and are irrespective of graft size or TH+ cell number.

Manual analysis of projection density revealed greater neurite outgrowth into both the lateral and medial striatum for d18 grafts, compared to d25 grafts (**Figure 6G**; DIV. F_1,11_= 12.12 and 37.54, respectively, ps≤0.005). Overall an approximately 2.3-fold increase in neurite outgrowth was evident in the medial striatum, relative to the lateral striatum (**Figure 6G**; Striatal region. F_1,11_=287.83, p<0.001). Unbiased optical density analysis reflected this same pattern of innervation, with greater neurite outgrowth medial to the graft (**Figure 6H**; DIV. F_1,11_= 13.95 and 30.69 respectively, ps≤0.003).

Although differences were evident in neurite outgrowth from the graft into the surrounding host striatum, by 1400μm from the graft border no differences in the length or density of projections were observed between d18 and d25 grafts [**Figure 6A**; F_19,646_=13.22, p<0.001, 0-1300μms, ps<0.05; 1400-2000μm, ps=n.s.). Representative images of TH+ neurites are depicted in Figure **6B-E**.

Improvements in amphetamine-induced rotational bias correlated significantly with greater overall neurite outgrowth (**Figure 6I**; r= −0.754, p≤0.001), greater fiber projections into the lateral neostriatum (**Figure 6J**; r= −0.696, p≤0.001) and greater fiber outgrowth into the medial striatum (**Figure 6K**; r= −0.723, p≤0.001). This is further supported by correlation analysis of optical density measurements and 24 week amphetamine score, with reduced rotational bias associated with overall neurite outgrowth (**Supplemental Figure 2C**; r= - 0.388, p=0.021), lateral fiber outgrowth (**Supplemental Figure 2D;** r= −0.401, p=0.017) and medial fiber outgrowth (**Supplemental Figure 2E;** r= −0.521, p=0.001).

### Gene expression analysis identified signatures of developing DA neuron precursors

RNA sequencing was used to create phenotypic profiles of the cell cultures at each stage of DA neuron precursor development and to understand the trajectory of differentiation. Both cell lines were analyzed at the two stages used for efficacy assessment (d18 and d25). To investigate the changes over time, the cultures were also analyzed at day 13 of differentiation, and data from two undifferentiated iPSC lines from the publicly available HipSci database (https://www.phe-culturecollections.org.uk/products/celllines/hipsci/index.jsp)were included as day 0 profiles for comparison.

Principal component analysis (PCA) of the time course of differentiation (d13, d18, d25), including two unrelated iPSC lines for comparison (d0), showed no overlap among the time points; PC1 essentially separated the undifferentiated iPSC lines from the others and PC2 resolved the later timepoints in a consistent rank order. (**Figure 7A**). When only the three differentiation phases (d13, 18, 25), are compared, PC1 separated the three stages of differentiation for both cell lines, and PC2 distinguished the two cell lines that had similar but not identical profiles (**Figure 7B**). The factor loading of the top- and bottom-ranked genes on these PC axes are listed in **Supplemental Table S1**.

**Figure 7.**
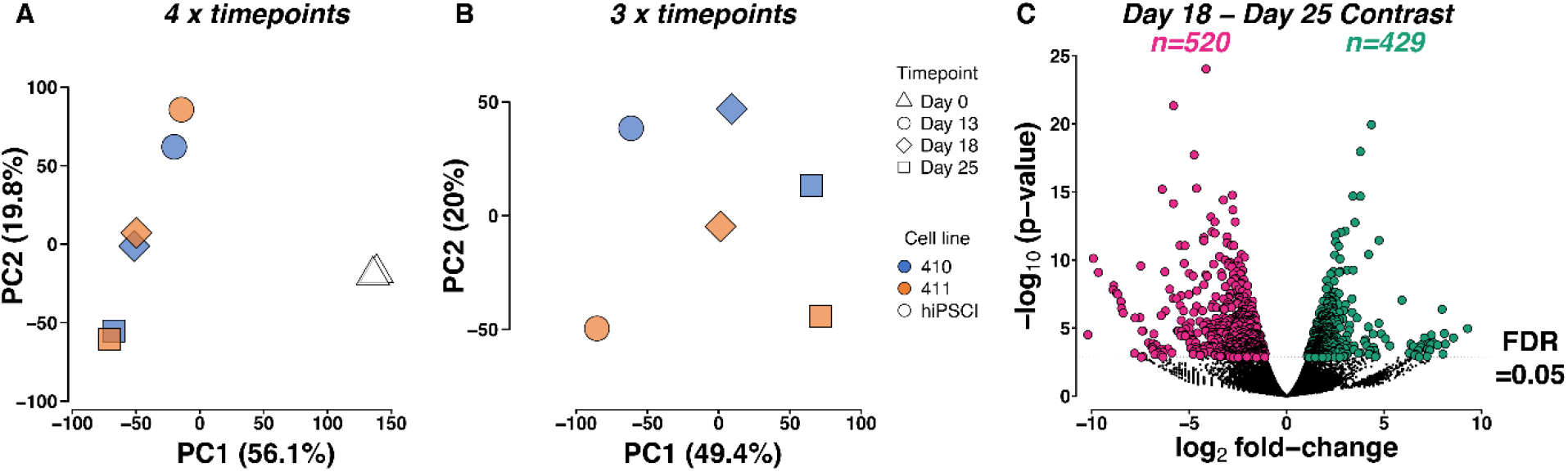
RNAseq-Global transcriptome variation and differential expression across differentiation timepoints. Principal component one (PC1) explained 56.1% of the variance in transcriptome expression in the four-time point data set (**A**) and largely separated the iPSCs (triangles: day 0) from the differentiated dopaminergic precursors (circles, diamonds and squares representing day 13, day18 and day 25, respectively). The second axis (PC2) delineated the DA precursors in a rank order corresponding with their development duration. The three-time point analysis (differentiation stages only) showed a strong signal of development stage along PC1 (49.4% of transcriptome variance) and PC2 separated the two cell lines (410: blue and 411: orange) within a time point (**B**). (**C**) Volcano plot. A total of 520 genes were up-regulated at day 25 relative to day 18 (pink). 429 genes were down-regulated between day 18 and day 25 (green) (FDR<0.05).

Differential expression (DE) analysis comparing the two stages used in the animal studies, (d18 and d25) showed 949 genes that were significantly (FDR<0.05) differentially expressed. Of these, 520 genes (FDR<0.05) were up-regulated in d25 compared to d18 and 429 genes (FDR<0.05) were down-regulated (**Supplemental Table S2**: genes with negative log2fold change have higher expression at d25 relative to d18). **Figure 7C** illustrates the transcriptome contrast between d18 and d25.

The key processes active in d13, d18 and d25 cultures are shown in **Supplemental Figure 3**. As expected, cell cycle genes were more enriched in the earlier cultures compared to d25; these include *CCNB2, AURKB, PTTG1* and *TOP2A*. Also, transcription factors associated with neural precursors (*NEUROG2, HES1, HES5, REST*) tended to be higher at d13 and d18 compared to d25, while transcription factors associated with specific dopaminergic neurogenesis, such as *LMX1A* and *NR4A2* (*NURR1*) were enriched in d25 cultures. Genes associated with developing neural precursors (*NES, SOX2, SOX9, RFX4*) were more highly expressed at the earlier stages, while genes expressed in dopaminergic neurons were more highly expressed at d25, including *TH, DDC, PBX1, PITX3* and *RET*. Some genes associated with astrocytes, such as *GFAP* and *SLC1A*, were identified at the two later stages, as were markers of oligodendrocytes (*OLIG2*), and, as reported previously (Tiklová *et al*., 2020), genes associated with vascular leptomeningeal cells (*COL1A1* and *COL1A2*).

Transcription binding site motifs for the transcription factors *E2F4, FOXM1, SIN3A* and *NFYA* were enriched in genes expressed at higher levels in day 18 cultures compared to the later stage (d25). Among the genes up-regulated at the later stage (d25) relative to d18, there was a strong enrichment for transcription factor binding site motifs for *REST, SUZ12, EZH2* and *SMAD4* (**Supplemental Table S3**).

Gene Ontology (GO) analysis indicated that the main gene set that was up-regulated at d18 relative to d25 was largely associated with proliferation. On d25, the up-regulated genes had an overwhelming signal of synapse-related ontologies (**Supplemental Table S4**).

Some genes that were specifically upregulated in d18 cells are associated with neurite outgrowth, including *LIN28A, FLRT3, and ITGA5. LIN28A* (log2FC=3.8) codes for a post-transcriptional regulator of miRNAs associated with embryogenesis and has been linked to axonal regeneration (Bhuiyan *et al*., 2013; Wang *et al*., 2018; Xia, Teotia and Ahmad, 2018; Nathan *et al*., 2020). Overexpression of *LIN28A* in DA neurons has been reported to increase dendrite length, graft volume, and TH+ content, and enhance functional recovery post-transplantation (Rhee *et al*., 2016). *FLRT3*, which was upregulated in d18 cells, is implicated in neurite outgrowth and has been identified as a positive regulator of FGF signaling and cell adhesion (Robinson *et al*., 2004; Tsuji *et al*., 2004). *FLRT3* codes for a co-receptor for Robo1; the attractive response to the guidance cue Netrin1 has been shown to be controlled by Slit/Robo1 signaling and by *FLRT3* (Tsuji *et al*., 2004; Leyva-Díaz *et al*., 2014). Thus, the expression of *FLRT3* may promote neurite outgrowth from the grafted d18 precursors. *ITGA5* codes for subunit alpha 5 in the integrin alpha chain family (Integrin α5β1), which has been identified as having a role in specific dopaminergic neuron outgrowth onto striatal neurons (Izumi *et al*., 2017).

## Discussion

Success of regenerative therapies using pluripotent stem cell-derived cell types is dependent on the demonstration of their functional efficacy, and the maturation stage and plasticity of the cultures is key to achieve this. There is not yet an established path to production of hPSC-derived cells that meets both of these criteria. Strategies differ depending on the disease and organ characteristics; for approaches that can use well-differentiated derivatives, such as transplant of iPSC-derived sheets of retinal pigment epithelium for age-related macular degeneration (Kamao *et al*., 2014), the final product can be assessed for correct identity and functionality pre-transplantation. However, for many applications, cells must be at a pre-differentiated precursor phase; for hPSC-derived dopamine neuron replacement, extensive independent studies indicate that they need to be far enough along the path of differentiation that they are committed to a narrow range of fates (“determined”-stage cells) but not so differentiated that they have extended neurites and have begun to make synapses. Mature neurons do not survive the dissociation process required to distribute the cells in the transplant site (Kim, Koo and Studer, 2020).

To assess reproducibility and efficacy of dopamine neuron precursors, we used iPSCs derived from two individuals with Parkinson’s disease to gain insight into the factors that affect successful engraftment and behavioral recovery in a rat model of PD. We found that the two iPSC lines differentiated along a similar time course and were capable of maturation *in vitro* into neurons that were electrophysiologically similar, expressed the characteristic markers, and synthesized and released dopamine. Dopaminergic neuron precursors derived from both cell lines engrafted well and differentiated *in vivo* into TH-positive cells with similar efficacy. This shows that it is feasible to produce very similar cell types from multiple iPSC lines.

However, there was a striking difference in the functionality of the grafts that was dependent not on the cell line but on the extent that the cells were differentiated prior to transplantation. Both cell lines alleviated the rats’ functional deficits when they were cultured for 18 days before grafting, but in stark contrast, cells from both cell lines, when cultured for a week longer, 25 days, did not restore the behavioral deficits even though they engrafted well and expressed TH, AADC and GIRK2, typical markers of dopamine neurons.

The likely underlying cause of the differences in efficacy between the two timepoints was identified through histological analysis. The grafts were variable in volume, density and quantity of iPSC-derived dopaminergic neurons, but these characteristics did not correlate with the functional efficacy. Instead, histological analysis revealed distinct differences in the extent of neurite outgrowth into the host brain between the two timepoints. Unlike the d25 grafts, the d18 grafts had extensive outgrowth with dominant innervation towards the medial subregion of the striatal nucleus. The dense innervation overall correlates well with functional recovery.

Our results indicate that it is the propensity for neurite outgrowth into the host striatal nucleus that underpins behavioral recovery, while the survival of dopaminergic neurons in grafts is not sufficient to ensure efficacy. Thus, our findings identify graft-host connectivity as a key feature of graft efficacy.

Gene expression profiling has been used for many years to provide unbiased characteristics of cell phenotypes and to predict the differentiation abilities of pluripotent stem cells (Müller *et al*., 2011; Allison *et al*., 2018), and it is likely to prove useful for prediction of other qualities, such as post-transplantation characteristics. This unbiased approach provides an opportunity to understand what genes expressed at the efficacious stage (d18) may be drivers of neurite outgrowth and genes at the ineffective stage (d25) whose differential expression might be used as markers of reduced outgrowth potential. In general, our gene expression analysis suggested that the cultures that were functionally effective were poised to begin to mature, but had not quite initiated the processes leading to functional neurons. One gene that is of interest is *REST* (RE1 silencing transcription factor), which codes for a transcription factor that acts as a repressor of genes involved in neural maturation; its expression is thought to allow a pool of neural precursors to accumulate during processes of neural differentiation in embryogenesis (Hwang and Zukin, 2018). In our cultures, *REST* was detected in the earlier cell stages but decreased considerably at d25; in addition, at day 25, genes with REST transcription factor binding motifs were upregulated, which is consistent with removal of REST suppression. *REST* expression and expression of the regulated genes could prove to be useful markers to assess an optimal stage of transplantation of future cell therapy products.

In conclusion, our study is unique in that we have distinguished two maturation stages that are both capable of surviving long-term *in vivo* and producing dopamine neuron-rich grafts in a rat model of PD, but observed that only one of the stages restored motor function. The effective cells behaved differently in the host brain, producing extensive neurite outgrowth. It is important to note that different differentiation protocols result in distinct maturation timescales, so the specific number of days in culture that produces optimal *in vivo* transplants will differ according to the protocol (Kirkeby *et al*., 2012; Qiu *et al*., 2017; Song *et al*., 2020). Our work does suggest, however, that it is likely that the optimal time of differentiation before transplantation may be identifiable for all protocols. Phenotyping cultures by gene expression analysis offers insights into the characteristics that distinguish cells that develop into effective grafts from those that fail, and may be useful for discovering markers that can predict the success of cultures. Gene expression profiles may eventually be used to identify an optimized product to ensure extensive neurite outgrowth and consequent functional recovery. This is especially important for autologous cell replacement therapy for PD (Loring, 2018), for which effective cell types must be derived reproducibly from each graft recipient.

## Methods

### Experimental Overview

Two dopamine precursor cell preparations derived from different PD patients (lines 410 and 411) were cryopreserved at either day 18 or day 25 of the differentiation protocol. Replicate cultures were further differentiated and analyzed by immunocytochemistry, HPLC and electrophysiology assays. Lister-hooded rats were kept either as naïve controls or received 6-OHDA lesions to the medial forebrain bundle (MFB) (**Figure 4**). At 3− and 4− weeks post-lesion, rats were tested with the amphetamine-induced rotation test and were distributed into matched groups based on net ipsilateral rotations. At 5 weeks post-lesion, subsets of lesioned rats were either maintained as lesion-only controls or received intrastriatal transplants of one of the following preparations: iPSC line 410 differentiated to d18 (410− d18), 410 at d25 (410-d25), line 411 at d18 (411-d18) and 411 at d25 (411-d25) cells. Rats were maintained on daily cyclosporine A immunosuppression at 10mg/kg and underwent amphetamine rotation assays every 4 weeks post-lesion. At 24-weeks post-graft, rats underwent gait analysis and an apomorphine rotation test. All rats were subsequently perfused with fixative, and the brain tissue harvested for immunohistochemical analysis.

### Cell culture

With informed consent and Institutional Review Board approval (Scripps IRB HSC-08-5109) dermal punch biopsies (3mm) were obtained from two individuals diagnosed with idiopathic PD: donor 410− Male, 62 years, Caucasian; and donor 411− Male, 51 years, Caucasian. Dermal fibroblasts were isolated as described (Glenn *et al*., 2012) and reprogrammed using the Sendai CytoTune-iPS Reprogramming Kit (Thermo Fisher). Multiple iPSC clones from each line were isolated, expanded and banked as previously described (Boland *et al*., 2017).

#### Differentiation

iPSCs were differentiated on Geltrex (Life Technologies, 1:200 dilutions) using a modified version of a previously published dual-SMAD inhibition protocol (Kriks *et al*., 2011). iPSCs were dissociated with Accutase (Gibco) and seeded as single cells at a concentration of 200K cells/cm^2^ in maintenance medium (Essential 8 medium, Thermo Fisher) supplemented with a rho kinase inhibitor (Stemgent, 04-0012-02, 1μM) before switching to differentiation medium 24 hours later. Differentiation medium consisted of 1:1 mix of DMEM/F-12 and Neurobasal medium containing 1 x N2/B27, Glutamax and MEM-NEAA (all from Thermo Fisher). Differentiation medium contained varying amounts of KnockOut Serum Replacement (Thermo Fisher) starting at 5% on the first 2 days of differentiation, decreasing to 2% through day 10 of differentiation. The following were added to the differentiation medium to induce floor plate precursor differentiation: LDN 193189 (days 1-13; 100nM, Stemgent), SB431542 (days 1-5, 2μM, Tocris), CHIR99021 (days 3-13, 2μM, Stemgent), Purmorphamine (days 2-7, 2μM, Calbiochem), sonic hedgehog C25II (days 2-7, 100ng/mL, R&D Systems). After day 13 of differentiation, basal medium was switched to Neurobasal medium containing 1x N2/B27, Glutamax, and MEM-NEAA supplemented with BDNF (20ng/mL, R&D systems), GDNF (20ng/mL, Preprotech), ascorbic acid (0.2mM, Sigma-Aldrich), dBcAMP (0.5mM, Sigma-Aldrich), TGFB3 (1ng/mL, R&D Systems) and DAPT (10μM, Tocris). On day 16 of differentiation, cells were passaged using Dispase/collagenase (Roche) and DNase (Worthington Biomedical) and reseeded as at 1:2 passage ratio on poly-l-ornithine- (Sigma), laminin- (Roche) and fibronectin- (Sigma) coated dishes in medium containing rho kinase inhibitor. For cells transplanted on d18 of differentiation, cultures were treated with Accutase and a single-cell suspension was cryopreserved in CryoStor CS10 cryopreservation medium (Stem Technologies) according to manufacturer’s instructions. For day 25 transplantation, parallel cultures were passaged at day 20 using Accutase, reseeded at a 1:1 ratio in poly-l-ornithine (Sigma), laminin (Roche) and Fibronectin (Sigma) coated dishes and cultured to day 25, when they were dissociated with Accutase and single-cell suspensions were cryopreserved. Parallel cultures of each line were allowed to mature further for about 50-60 days. Laminin was supplemented into the medium once a week at a concentration of 1μg/ml to maintain attachment to the surface.

### Immunocytochemistry of dopamine neurons

After 80 days of differentiation, cells were fixed with 4% paraformaldehyde (Electron Microscopy Sciences), washed and blocked in PBS containing 10% Triton x-100 (MP Biomedicals), 5% donkey serum (Sigma) and 0.2% bovine albumin (Fisher) for 30 minutes at room temperature. Anti-tyrosine hydroxylase (Millipore AB 152, 1:500) was added to the cells for 1 hour at room temperature or overnight at 4°C. Cells were washed in blocking buffer with Hoechst nuclear stain (1:10,000) and mounted with Prolong Gold Antifade (Thermo Fisher).

### HPLC

Mature neuronal cultures (88-89 days in vitro) were depolarized with potassium chloride solution to elicit neurotransmitter release. Culture medium was collected and HPLC analysis for dopamine, serotonin and norepinephrine was conducted (See Supplemental Methods for detailed description of the methodology).

### In vitro electrophysiology

Electrophysiological recordings were performed *in vitro*, in current-clamp mode, in order to characterize elements of the action potentials in spontaneously firing neurons. (See Supplemental Methods for detailed description of the methodology).

### Gene Expression analysis of cell cultures

A detailed description for RNAseq pre-processing and analyses can be found in Supplemental Methods. Briefly, we prepared total RNA libraries for paired-end sequencing from six cell culture samples: d13 (n=2), d18 (n=2) and d25 (n=2). For all three timepoints one biological sample was from the 410 cell line and the other was from the 411 cell line. In addition, we included two RNAseq libraries from the HiPSCl collection (https://www.phe-culturecollections.org.uk/products/celllines/hipsci/index.jsp) as examples of undifferentiated pluripotent cells for which RNAseq data were publicly available (full detailed available in Supplemental Methods).

The differential expression (DE) analysis reported in the main text compared the focal timepoints that were transplanted into rats: d18 and d25 cells. We also performed principal component analyses (PCAs) to understand global changes in gene expression across the differentiation timepoints. The output from the DE test was used to conduct enrichment tests across gene ontologies (GO) and transcription factor binding site (TFBS) motifs.

### Animals

Sixty-five adult female Lister-Hooded rats were used, weighing 200-250g at the time of the first surgical procedure-Animals were housed under standard conditions with free access to food and water. All experiments were conducted in accordance with the requirements of the UK Animal Scientific Procedures Act, 1986, under Home Office PIL P49E8C976. All animals received daily administration of 10mg/kg cyclosporine post grafting surgery for the duration of the experiment.

#### Group inclusion

Behavioral and histological analysis was performed on fifty-one animals; groups sizes were: Control (n=9), Lesion only (n=7), Lesion with transplants, 410-d18 (n=8), 410-d25 (n=8), 411-d18 (n=12), 411-d25 (n=7). Fourteen animals were excluded from final data analysis; these were distributed across all grafted and non-grafted groups (**Supplemental Figure 4**). Seven animals were culled before the final post-graft time point due to health concerns over prolonged cyclosporine treatment. An additional seven were excluded after initial histological observation because they had either: no surviving graft (n=2), a graft with evidence of unusual proliferation (n=1), or because the grafts were too small to result in behavioral recovery (n=4). Small grafts were excluded because studies using both human fetal VM (Brundin *et al*., 1985; Rath *et al*., 2013) and hPSC-derived dopamine neurons (Grealish *et al*., 2014) have shown that at least 500 dopamine neurons per graft are required to induce significant behavioral recovery. Excluding small grafts ensures that analyses were specific to the relationship between functional recovery and fiber outgrowth without skewing the dataset by inclusion of grafts likely to be too small to support behavioral recovery.

#### General surgery

For all surgeries, anesthesia was first induced with 5% isoflurane in O_2_ (1l/min) and maintained with 1.5-3% isoflourane in O_2_ (0.8L/min) and NO (0.4L/min). Animals were placed in a stereotaxic frame (Kopf 900, Germany) with the nose bar set to −3.2 mm below the interaural line. Anterior-posterior and lateral coordinates were taken from bregma and ventral coordinates from dura.

#### Lesion surgery

Unilateral dopamine lesions were induced by the administration of 6− hydroxydopamine hydrobromide (Sigma, H116-5MG) into the MFB at the following coordinates: a=-4.0, L=-1.3, V=-7.0. Toxin was used at a free base weight concentration of 3ug/μl and delivered via cannula at a rate of 1ul/min over 3 min. Animals received a 5 ml subcutaneous infusion of glucose-saline and 1mg/kg Metacam as analgesia.

#### Transplantation surgery

Animals received a unilateral transplant of cells into the lesioned striatum. iPSCs were thawed in a ThawSTAR^®^ system (BioCision) and a cell suspension of 250,000cells/μl made in HBSS with Pulmozyme DNase. The cells were kept on ice until grafting and discarded for a fresh cell suspension after 4 hours. Grafted rats received a total of 500,000 cells in 2μL at stereotaxic coordinates: A=+0.6, L=-3.0, V=-4 and −5 (with nose bar set at −3.2mm).

### Drug-induced Rotations

All rotations were assessed using an automated rotometer system, with Rotorat software (Med Associates Inc. product: SOF-801).

#### Amphetamine-induced rotations

Rats received intraperitoneal administration of methamphetamine (Sigma, M8750-5G), at a dose of 2.5mg/kg in sterile saline. Amphetamine-induced rotation assays were carried out at 4-weeks post lesion as a baseline and then every 4 weeks post graft until end of experiment (24 weeks post-graft). Net rotation scores were calculated by subtracting contralateral from ipsilateral turns for each 60-second block and averaging over the total 90 min trial time. Animals achieving amphetamine-induced rotation scores of <6 rotations/min were excluded from the experiment (Björklund and Dunnett, 2019).

#### Apomorphine-induced rotations

At 24 weeks post-graft, rats were injected with a subcutaneous administration of apomorphine (Sigma, A4393-250MG), at a dose of 0.05mg/kg in sterile saline. Ipsilateral vs contralateral rotations scores for each 60 second block were expressed as total net rotation scores over a total trial time of 40 minutes.

#### Gait Analysis

At 24 weeks post-graft, animals were trained to traverse a paper-lined corridor. Animals completing the task in ≤3 seconds had their forepaws painted blue and hindpaws painted red and were allowed to traverse the corridor again. The paper with painted paw prints was saved from each run. The stride length was analyzed by measuring the distance from the centers of each paw to corresponding paw for each stride, with global stride length being an average of all measurements, (from both; contralateral and ipsilateral sides, and forepaw and hindpaw). An average of 3 stride lengths were taken (Dunnett and Brooks, 2018).

### Perfusion and Tissue Preparation

Animals received an intraperitoneal overdose of pentobarbital, Euthatal (Merial, MBEUT01) and were pericardially perfused with ice cold PBS and 250ml of 4% paraformaldehyde (PFA), before being post-fixed in 4% PFA for 4 hours. Brains were saturated in 25% sucrose in PBS until equilibrized before being cut on a freezing, sledge microtome at 30μm.

### Immunohistochemistry and microscopy

Immunohistochemical staining was carried out on free floating sections. Sections were washed and maintained in TRIS buffered saline (TBS, pH-7.4). Endogenous peroxidase activity was quenched by 5-minute incubation in methanol (10%) and hydrogen peroxide (3%) and washed in TBS. Non-specific binding sites were blocked with 1-hour incubation in 3% horse or goat serum. Sections were then incubated overnight in the primary antibody at the required concentration (anti-TH (Millipore; 1:2000); anti-GIRK2 (Alomone, 1:500); anti-AADC (Abcam, 1:1000); anti-HuNu (Millipore, 1:1000). After washing, the complementary secondary antibody was added to the sections for 3 hours. Sections were then incubated for 2 hours with biotin-streptavidin kit (Vector labs, PK-6100). Specific staining was then visualized with either 3,3-diaminobenzidine (DAB) (Sigma-Aldrich, D12384) or Vector SG peroxidase (Vector laboratories, SK-4700). Stained sections were mounted on gelatin-coated slides and allowed to dry at room temperature for 24 hours before being dehydrated (70%, 95%, 100% and Xylene for 5 mins) and cover slipped with DPX.

### Stereology

2-dimensional quantification was carried out using an automated stage and stereology module NEWCAST TM from Visiopharm Integrator System software (Visiopharm, UK) on an Olympus Bx50 microscope (Olympus Optical Co., Ltd, Japan). The total number of HuNu+, TH+, GIRK2+ and AADC+ cells were estimated by counting all labelled cells within either a 1:6 or 1:12 series of grafted sections at 200x magnification.

### Analysis of graft projections

#### Projection Counting

Fiber projections were visualized using a magnification of 400x. A 100um boundary was placed 100um away from the graft border, any fibers crossing this boundary were manually counted. This was repeated at regular 100μm intervals moving away from the graft, until no fibers remained, or the boundary was no longer in the striatum. This was conducted in both medial and lateral directions from the graft. Where possible, five evenly spaced samples were taken from the brain section with the largest graft (Bagga, Dunnett and Fricker-Gates, 2008). To ensure independent analysis of fibers, if the five samples could not be spaced more than 300μm from each other for a small graft, additional analyses were undertaken on subsequent grafted sections. Data is presented as both; average fiber density per 100um distance from the graft (A) and the average fiber density of the neostriatal area around the graft.

#### Optical Density Analysis

To verify the patterns seen in manual projection counting and objective measurement of fiber density was taken from the portions of the striatum both lateral and medial to the graft. High resolution images of each TH stained, grafted section were taken using an automated stage on an Olympus Bx50 microscope. Optical density was calculated using ImageJ software and normalized to a control reading taken from the cortex to correct for background staining. Fiber density is presented as a change in optical density from the cortical control region.

#### Statistical analyses

Amphetamine-induced rotations scores were analyzed using a repeated measures ANOVA with factors of group (control, lesion, 410-d18, 411-d18, 410− d25 and 411-d25) and experimental timepoint (pre-lesion baseline and then 4-, 8-, 12-, 16-, 20− and 24 weeks post-graft). Apomorphine net rotations scores and gait analysis stride length were analyzed using a one-way ANOVA, with Tukey HSD test for post-hoc analyses. All stereological analysis was undertaken using a repeated measures ANOVA, with cell line (410 or 411) and cryopreservation timepoint (d18 and d25) as factors. Bonferroni’s post hoc test was used to analyse simple effects for all ANOVA interaction terms.

For the projection analysis, factors of cell line, DIV, direction (medial vs. lateral) were used. Data for correlational analysis were analyzed for normality using a Kolmogorov–Smirnov test. For non-parametric data (i.e., all projection data), Spearman’s test of correlation was used instead of Pearson’s coefficient. All phenotype statistical analysis was carried out using SPSS software, version 25.0 (IBM, Armonk, NY).

## Acknowledgements

MJL was supported by a Parkinson’s UK Senior Research Fellowship (F-1502); RH was supported by a grant from Summit for Stem Cell Foundation; JFL, HT, ABL, and RMW were supported by grants from Summit for Stem Cell Foundation and the California Institute for Regenerative Medicine (CIRM; CL1-00502, RT3-07655, GC1R-06673-A) JFL, HT, ABL, RMW and DGS were supported by CIRM (DISC2-09073); PPS and IVS were supported by NIH (DA046170, DA046204-04, DA043268). The authors wish to acknowledge our Scripps colleague, Loren (Larry) H. Parsons (1964-2016) for the use of his laboratory for HPLC analysis.

## Data Availability Statement

Raw RNA-seq reads generated in this study are available from the NCBI Gene Expression Omnibus (GEO) data repository (https://www.ncbi.nlm.nih.gov/geo/) under project accession: GSE186243.

## Supplemental Figures and Tables

**Supplemental Figure 1:**
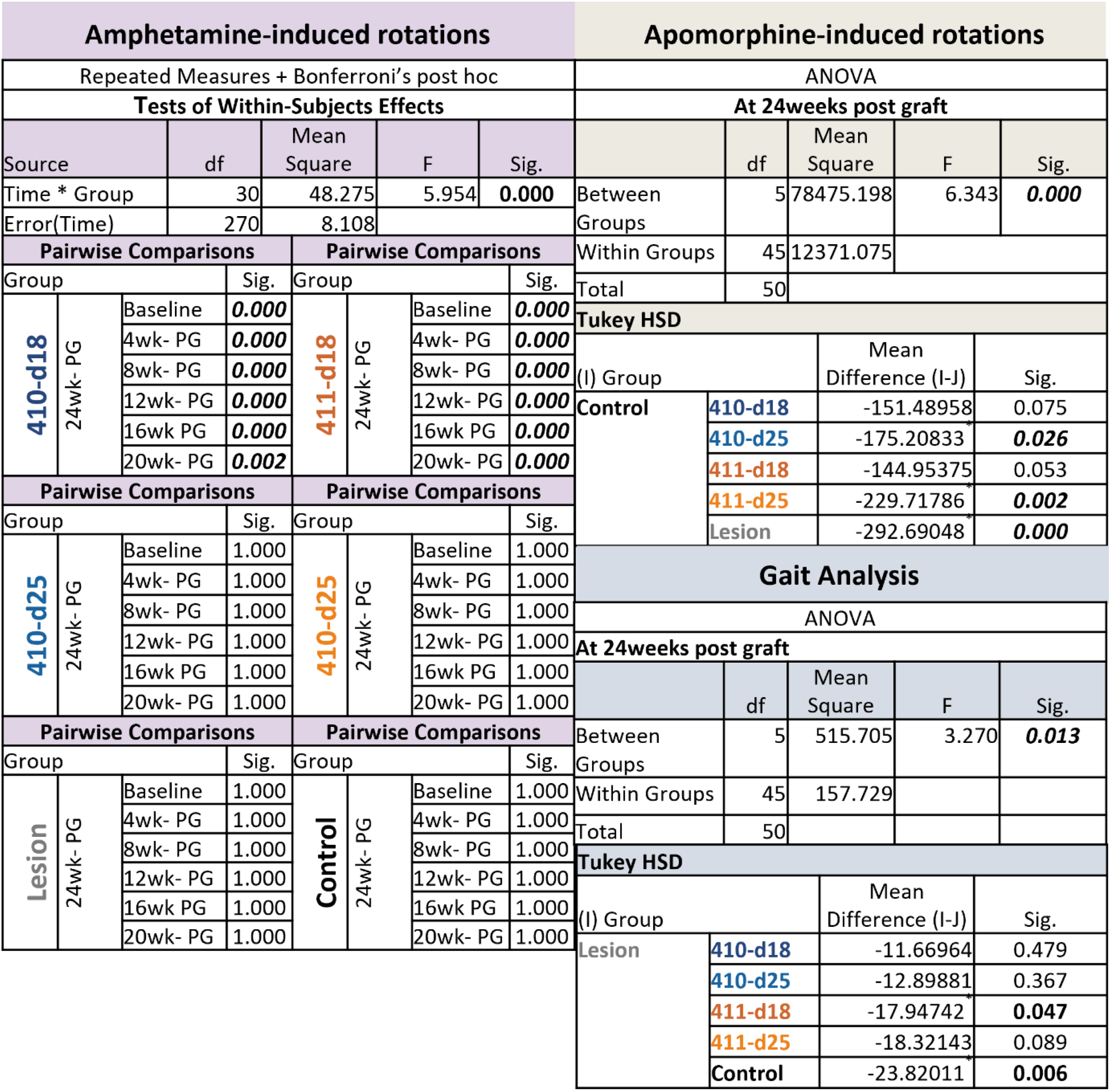
ANOVA table associated with the statistical analysis of amphetamine-induced rotation data, showing a summary of F- and P-values for statistical tests performed (repeated measures ANOVA with Group and Time as factors, and Bonferroni’s post hoc test). Oneway-ANOVA output for both apomorphine-induced rotations and gait analysis datasets, with Tukey’s post-hoc analysis. Significance, set at p<0.05, is denoted in bold.

**Supplemental Figure 2:**
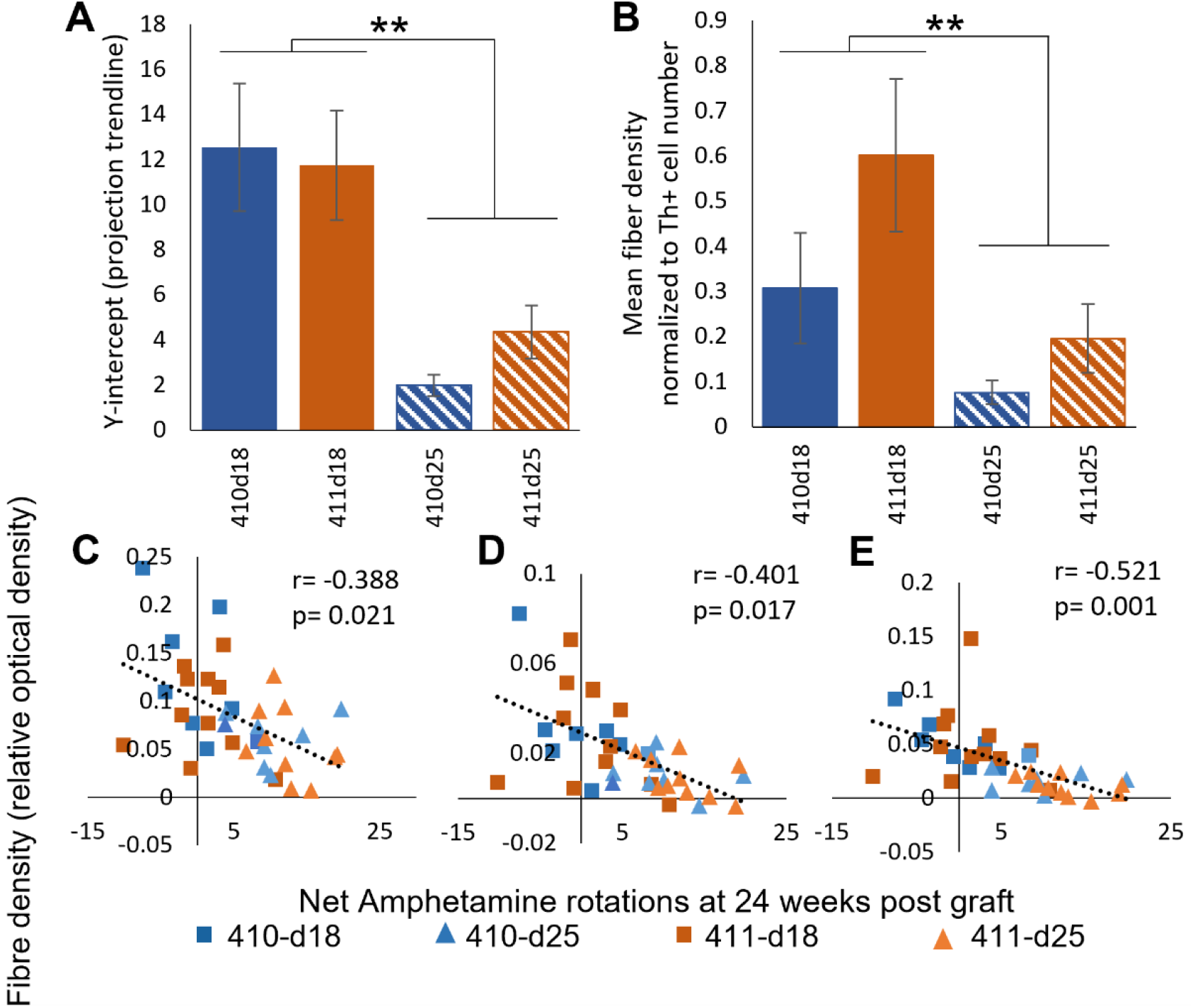
Additional analysis of the pattern of neurite outgrowth. (A) Analysis of the y-intercept data was undertaken and revealed significantly higher y-intercepts in the two d18 graft groups, as compared to the two d25 graft groups, indicating more projections overall in the d18 grafts. (B) Normalization of the neurite outgrowth data to the total number of TH+ cells was undertaken to eliminate the impact of any differences in graft size. The data reveal significantly more neurite outgrowth in both day18 graft groups, as compared to both d25 graft groups. (C-E) Correlational analysis of the optical density projection data and net amphetamine rotations at 24 weeks post-graft revealed a relationship between more (C) overall outgrowth, more (D) lateral outgrowth and more medial outgrowth and better recovery on the amphetamine-induced rotation test.

**Supplemental Figure 3:**
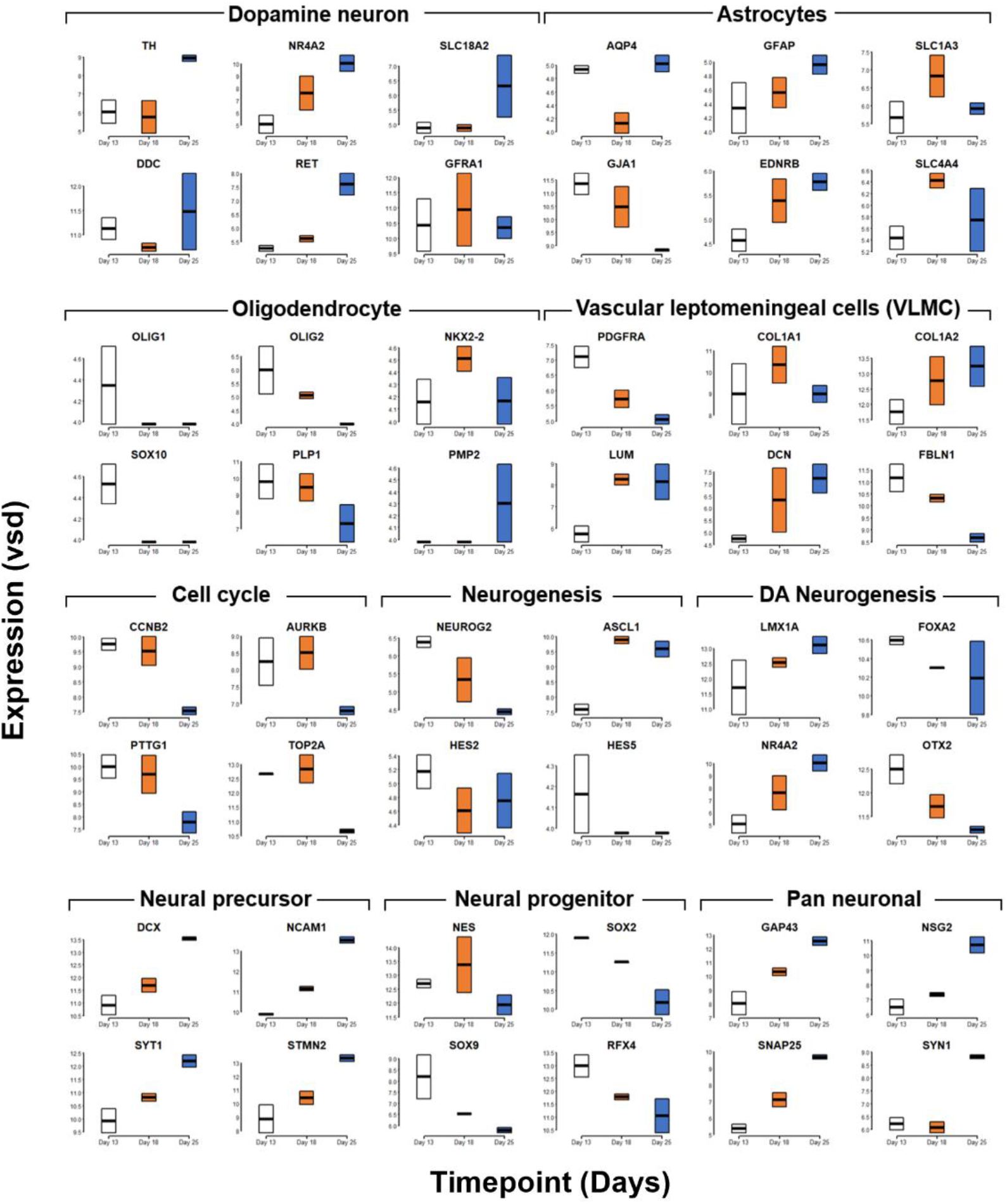
RNAseq data for cell preparations harvested at d13, d18 and d25, with genes grouped according to association with cell type or stage. Expression values are displayed as variance stabilized data (vsd), calculated using the vst command in the DESeq2 package (Love, Huber and Anders, 2014).

**Supplemental Figure 4:**
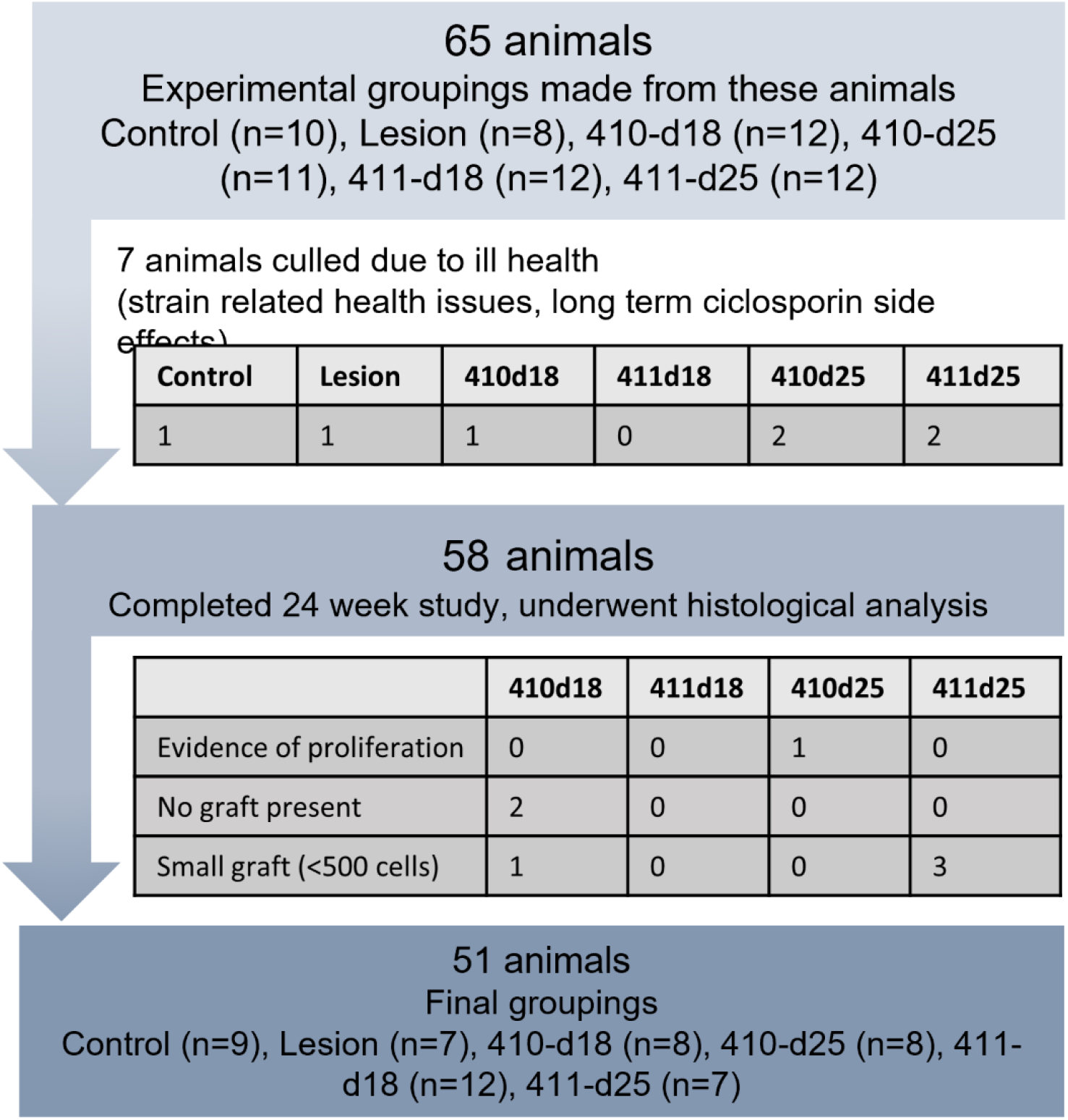
Schematic depicting the exclusion of rats from the study, detailing the stage of removal and the rationale for removal.

**Supplemental Table S1:** Principal component analysis. The top- and bottom-ranked factor loadings for PC1 and PC2 are shown for the analyses illustrated in **Fig 7A and B**. Rank corresponds with a sorted list (largest value → smallest value). In Fig 7A, PC1 largely distinguished the iPSC samples from the Day 13, Day 18, and Day 25 DA neurons. PC2 largely delineated the differentiation timepoints. In Fig 7B, PC1 largely corresponded with differentiation time, while PC2 resolved the between-sample (within-timepoint) differences.

**Supplemental Table 2:** Differential expression analysis comparing the two stages used in the animal studies (d18 and d25). 949 genes significantly (FDR<0.05) differentially expressed. 520 genes (FDR<0.05) up-regulated in d25 compared to d18 and 429 genes (FDR<0.05) down-regulated. Genes with negative log_2_fold change have higher expression at d25 relative to d18. Column labels: genename = Ensemble gene name; ENSG = stable Ensemble gene number identifier; logFC = log_2_ fold change; logCPM = log of counts per million; LR = likelihood ratio; P-value = unadjusted p-value; FDR = false discovery rate (Benjamini-Hochberg method)(Benjamini and Hochberg, 1995).

**Supplemental Table 3:** Transcription binding site motif enrichment in genes upregulated on d18 or d25.

**Supplemental Table 4:** Gene ontology (GO) terms for genes differentially expressed in d18 vs. d25.

