## Supplemental Methods for "Functional Recovery from Human Induced Pluripotent Stem Cell-Derived Dopamine Neuron Grafts is Dependent on Neurite Outgrowth"

Hills *et al*.

**HPLC**

Mature neuronal cultures (88-89 days *in vitro)* were depolarized with potassium chloride solution to elicit neurotransmitter release. Culture medium was aspirated and cells were washed with Krebs Ring Buffer (KRB, Sigma). Fresh KRB was added back to the cells and a baseline sample was collected 15 minutes after addition. KRB was aspirated and replaced with KRB containing 56 mM KCl to depolarize the cells. Supernatant was collected 15 minutes later and immediately frozen on dry ice.

For HPLC analysis, dopamine (DA-HCl, MW 189.64), serotonin (5HT-creatinine sulfate, MW 387.41), and norepinephrine (NE-Bitartrate, MW 319.26), were obtained from Sigma Aldrich, USA and prepared into a 40 uM standard reference solution in Milli-Q purified water. The standard solution was serially diluted into 5, 2.5, 1.25, and 0.625 nanomolar concentrations that were used to generate a standard curve. 5 uL of sample and standard were analyzed using an Eicom HTEC-500 system. Mobile phase was prepared using 700 ml Milli-Q water with 5.18 g ammonium acetate (Sigma A7330, MW 77.08), 7.18g sodium sulfate (Sigma E1644, MW 142.04), 164 microliters acetic acid (Sigma A6283 MW 60.05), 300 ml methanol (Fisher 60-048-242), and 1 ml of a 50 mg/mL EDTA solution (EDTA-2NA, Sigma S9627, MW 372.24). The HPLC system was set to a flow rate of 250 μl/min, temperature of 35° F, time constant of 3.0, and E Potential of 450mV, with an EicomPAK CAX (2mm x 200mm) column. The resulting peak areas were measured using Agilent Chemstation software.

**Electrophysiology**

Electrophysiological recordings were performed in a small-volume recording chamber providing laminar solution flow (Warner Instruments, Hamden, CT). A coverslip with the plated cells was placed in a recording chamber perfused with 32°C artificial cerebrospinal fluid (ACSF) with the following composition (mM): 124 NaCl, 2.5 KCl, 1.25 NaH2PO4, 26 NaHCO3, 2 CaCl2, 2 MgCl2 and 10 glucose (95% O2/5% CO2, all chemicals purchased from Sigma Aldrich, St. Louis, MO). Cells were visualized with infrared differential interference contrast (IR-DIC) using a water immersion objective (Nikon Instruments Inc., Melville, NY). Patch pipettes were pulled from borosilicate glass tubing (1.5 mm outer diameter, 0.3 mm wall thickness) on a horizontal puller (P-97; Sutter Instrument, Novato, CA). Pipettes were filled with solution containing 150 mM potassium gluconate, 5 mM NaCl, 5 mM HEPES, 0.5 mM EGTA, 0.1 mM CaCl2, 2 mM MgATP, 0.3 mM Na_3_GTP, 2 mM phosphocreatine at pH 7.3. Recordings made with Axopatch 700B amplifier (Molecular Devices, Sunnyvale, CA) were low-pass filtered at 3 kHz and digitized on-line at 20 kHz using National Instruments PCI-MIO-16-E4 multifunctional board controlled by the data acquisition software DASYLab 6.0 (Dasytec, Amherst, NH). The input resistance (Rm), the series resistance and the membrane capacitance (Cm) were evaluated by analyzing the steady-state and transient components of current responses to 5 mV voltage steps that were applied from the holding potential of −65 mV. Characteristics of the action potentials (APs) were evaluated using recordings in spontaneously firing neurons performed in current-clamp mode. The series resistance in the recordings included in the analysis did not exceed 25 MΩ.

**Gene expression analysis of cell cultures**

For each of the six samples, total RNA was extracted from approximately 1 million cells in culture using a *mir*VANA™ miRNA isolation kit (Invitrogen) following the manufacturer’s protocol. All samples achieved a minimum RNA Integrity Number (RIN) of 9.0 prior to sequencing. One hundred and fifty base pair (150bp) paired-end sequencing was performed on the Illumina HiSeq 2000 platform (Illumina, San Diego, CA).

Additional Day 0 (iPSC) RNA-seq samples for PCA reference*.* RNA-seq libraries from two iPSC lines were obtained from NCBI’s Sequence Read Archive (SRA: https://www.ncbi.nlm.nih.gov/sra) to enable a reference set for our differentiated cell lines in the PCA analysis. Both samples belonged to the same Human Induced Pluripotent Stem Cell Initiative (HIPSCI) collection BioProject ID (Kilpinen et al., 2017): PRJEB7388, study reference: ERP007111 and were analyzed using the same sequencing methodology as the present study (Illumina HiSeq 2000, paired-end). Sample 1: Accession: SAMEA2518322, Run ID: ERR914289, library name: 13086085. Sample 2: Accession: SAMEA2547619, Run ID: ERR914288, library name: 13086073. Read mapping of the iPSC samples was performed using an identical protocol to the six differentiated sample libraries (see Sequence library pre-processing, below).

Sequence library pre-processing. We used the nf-core-rnaseq v1.4.2 pipeline (<https://github.com/nf-core/rnaseq>) for sample preprocessing (Ewels et al., 2020).  Fastq files were interpreted and processed using the Salmon pseudo-aligner (salmon version 1.1.0)(Patro et al., 2017) using default parameters with no additional flags and the GENCODE human reference genome (release 32) as the genome index. Transcripts were aggregated (summed) for each gene so the sum of all transcripts per gene was the gene-level count. Count data were used for differential expression analyses.

RNA-seq library pre-filtering*.* To increase differential expression (DE) analysis sensitivity we performed pre-filtering on genes sensu (Anders and Huber, 2010). We removed the bottom 40% quantile of genes sorted by their row sums. This resulted in 32,836 genes remaining in the downstream DE analyses. Crucially, the pre-filtering step was informed only by the six libraries in the present study. The additional two iPSC (Day 0) libraries were trimmed of the same excluded gene set as identified in the libraries from the present study. The additional HIPSCI samples therefore did not influence the differential expression analyses in any way, since filtering was performed independently of them. Day 0 samples were not used in any DE analysis and serve as a reference for the PCA only.

Differential expression (DE) analysis*.* DE analyses were performed using edgeR (Robinson et al., 2009). The matrix design was a factor of day, resulting in three levels (Day 13, Day 18, Day 25), each containing two biological replicates. Dispersion estimates were calculated sequentially by estimateGLMCommonDisp, estimateGLMTrendedDisp, and estimateGLMTagwiseDisp methods. We constructed the differential expression contrast using the makeContrasts command and fit the gene-wise negative binomial generalized linear model using glmFit. Likelihood ratio tests were performed using the glmLRT test, while applying the Benjamini and Hochberg multiple test correction (Benjamini and Hochberg, 1995). A false discovery rate (FDR) < 0.05 was used as the nominal statistical threshold for DE tests.

Principal Components Analysis (PCA)*.* To understand the general trends across all pre-filtered transcripts in the dataset (n=32,836), we performed principal components analyses (PCA) using the prcomp method implemented in the stats R package. Raw counts were first normalized by their size factor and variance stabilized using the DESeq2 *vst* method (Love et al., 2014). We performed two analyses: (1) using the six libraries with two previously published induced pluripotent stem cell (iPSC) libraries representing Day 0, and (2) using only the six libraries from the present study. The purpose of the first analysis is to provide an outgroup reference of iPSCs, which are known to demonstrate a canonical pluripotent expression pattern (Allison et al., 2018).

Transcription factor binding site (TFBS) enrichment (cis-regulatory*).* We tested for over-represented transcription factor binding sites in the differentially expressed gene lists using the X2K webtool with default parameters (ChIP-seq experiments (CHEA) and ENCODE consensus background). If multiple entries were detected for a transcription factor, the version with the lowest hypergeometric test p-value was retained and the other removed from the data set for the purpose of figure clarity. Supplemental Table S3 contains the complete data set and all results are retained.

Kinase Enrichment Analysis (KEA*).* To test for signatures of protein phosphorylation pathways and potential kinome regulatory cascades, we conducted kinase enrichment analysis (KEA) (Chen et al., 2012; Lachmann and Ma’ayan, 2009), and implemented in the expression to kinase (X2K) web tool (amp.pharm.mssm.edu/X2K/). Default analysis parameters were used throughout. Kinase significance was calculated using hypergeometric tests against the background kinase and kinase-family distribution obtained from the KEA database.

Gene Ontology (GO) analysis*.* We tested for enriched gene ontologies in the gene sets that were: (a) up- or (b) down-regulated between the Day 18 and Day 25 samples. Gene ontologies were tested using the GOrilla program with an unranked test set compared to the background (Eden et al., 2009). The background data set was the total number of genes in the analysis (32,836) following good practice recommendations (Timmons et al., 2015). Of these, 20,158 genes were recognized, and 16,365 were associated with an ontology. We extracted all ontologies with an adjusted FDR q-value <0.05 in ‘Biological Process’, ‘Molecular Function’, and ‘Cellular Component’ categories. GO enrichment analyses are reported in Supplemental Table S4.

Behavior of canonical neuronal differentiation genes. We wanted to assess the behavior of genes in our dataset that are known to be associated with neuronal graft development. We used gene lists obtained from a recently published study on single cell transcriptomics in a model of Parkinson disease (Tiklová et al., 2020). These data are represented in Supplemental Figure S2.
