## Supplemental Table 1 for "Functional Recovery from Human Induced Pluripotent Stem Cell-Derived Dopamine Neuron Grafts is Dependent on Neurite Outgrowth"

**Supplemental Table S1**

|  |  | **Figure 7A** | | **Figure 7B** | |
| --- | --- | --- | --- | --- | --- |
|  | **Rank** | **PC1** | **PC2** | **PC1** | **PC2** |
| **Top-ranked (+ve)** | **1** | *TDGF1* | *CAPN6* | *POSTN* | *ADGRG7* |
|  | **2** | *ESRG* | *LIN28A* | *TPH1* | *AC018865.3* |
|  | **3** | *DPPA4* | *PRTG* | *AC010247.2* | *POTEF* |
|  | **4** | *POU5F1* | *TXNIP* | *POU2F2* | *TBC1D3C* |
|  | **5** | *L1TD1* | *DBX1* | *SLC1A2* | *CCDC144NL-AS1* |
|  | **6** | *MIR302CHG* | *COL22A1* | *CHGA* | *FEZF2* |
|  | **7** | *AL161431.1* | *NGFR* | *NR4A2* | *AP001347.1* |
|  | **8** | *LRRK1* | *P4HA1* | *PPP2R2B* | *PCAT7* |
|  | **9** | *HHLA1* | *FABP7* | *CDH20* | *WNT8B* |
|  | **10** | *VRTN* | *DOCK10* | *APC2* | *GDF7* |
|  | **11** | *ZSCAN10* | *BARHL2* | *PIEZO2* | *MAB21L2* |
|  | **12** | *SFRP2* | *RFX4* | *GLRA2* | *MECOM* |
|  | **13** | *LINC00678* | *LIX1* | *SYT13* | *TAFA5* |
|  | **14** | *LIN28A* | *PMEL* | *OLFM3* | *MTUS1* |
|  | **15** | *AC009055.2* | *SLC16A3* | *SV2B* | *SLITRK6* |
|  | **:** | **:** | **:** | **:** | **:** |
| **Bottom-ranked (-ve)** | **32,822** | *NTN1* | *NPTX1* | *TXNIP* | *C9orf64* |
|  | **32,823** | *ABCA8* | *CBLN1* | *LDHA* | *ZNF728* |
|  | **32,824** | *AUXG01000058.1* | *ESRP1* | *FABP7* | *CNTN4* |
|  | **32,825** | *FOXA1* | *NMNAT2* | *CRABP1* | *ZNF667* |
|  | **32,826** | *DMRTA2* | *PIEZO2* | *TPM2* | *SLC15A4* |
|  | **32,827** | *LRP2* | *CDH1* | *CACNA1G* | *ZNF667-AS1* |
|  | **32,828** | *MAP6* | *PPP2R2B* | *COL4A1* | *CHL1* |
|  | **32,829** | *LMX1B* | *SLC1A2* | *P4HA1* | *ZNF676* |
|  | **32,830** | *DDC* | *SYT13* | *ANKRD33B* | *ZFP28* |
|  | **32,831** | *RFX4* | *AC010247.2* | *SALL4* | *CHCHD2* |
|  | **32,832** | *PTPRO* | *JPH4* | *PRTG* | *ZNF736* |
|  | **32,833** | *KCNJ16* | *POU2F2* | *NGFR* | *ZNF208* |
|  | **32,834** | *LMX1A* | *TPH1* | *PMEL* | *TMEM132C* |
|  | **32,835** | *LINC00261* | *CHGA* | *CAPN6* | *PEG3* |
|  | **32,836** | *NNAT* | *POSTN* | *LIN28A* | *MEG3* |
